# Utilization of the genetically encoded calcium indicator Salsa6F in cardiac applications

**DOI:** 10.1101/2023.11.22.568284

**Authors:** Karla M. Márquez-Nogueras, Elisa Bovo, Quan Cao, Aleksey V. Zima, Ivana Y. Kuo

## Abstract

Calcium signaling is a critical process required for cellular mechanisms such as cardiac contractility. The inability of the cell to properly activate or regulate calcium signaling can lead to contractile dysfunction. In isolated cardiomyocytes, calcium signaling has been primarily studied using calcium fluorescent dyes, however these dyes have limited applicability to whole organs. Here, we crossed the Salsa6f mouse which expresses a genetically encoded ratiometric cytosolic calcium indicator with a cardiomyocyte specific inducible cre to temporally-induce expression and studied cytosolic calcium transients in isolated cardiomyocytes and modified Langendorff heart preparations. Isolated cardiomyocytes expressing Salsa6f or Fluo-4AM loaded were compared. We also crossed the Salsa6f mouse with a floxed Polycystin 2 (PC2) mouse to test the feasibility of using the Salsa6f mouse to measure calcium transients in PC2 heterozygous or homozygous knock out mice. Although there are caveats in the applicability of the Salsa6f mouse, there are clear advantages to using the Salsa6f mouse to measure whole heart calcium signals.

## 1. Introduction

Calcium is a secondary messenger used by cells to regulate and activate important cellular processes in a variety of organs including the heart. Cardiomyocyte contractility is governed by increase of cytosolic calcium due to the coupling of calcium influx in the plasma membrane and release from the sarcoplasmic reticulum (SR) [1]. Dysregulation of calcium homeostasis, due to mutations in calcium handling molecules or uncoupling of calcium handling, can lead to maladaptive responses resulting in heart failure [2]. Thus, the measurement of calcium signaling within cardiomyocytes is a mainstay of cardiac research. Calcium transients in ventricular or atrial cardiomyocytes have been primarily characterized through the loading of membrane permeant calcium dyes like Fura-2AM and Fluo-4AM and their derivatives. Due to the movement artefacts of a contracting cardiomyocyte, high-affinity ratiometric indicators such as Fura-2AM and indo-1, are commonly used in simultaneous calcium-contractile measurements. In cardiomyocytes when the contractile apparatus is pharmacologically inhibited, single wavelength dyes such as the low affinity indicator Fluo-4AM have been widely used, especially in applications with line-scan imaging [3, 4].

Although there are advantages to understanding calcium dynamics within a single cell, there are limitations in extrapolating isolated cardiomyocyte behavior to whole heart behavior. Indeed, a single cardiac cycle requires coordinated communication between the atria and ventricle starting with the electrical impulse that leads the contraction and ends with relaxation [5]. Thus, tools that enable the measurement of calcium in whole organ and how this is altered in disease states becomes relevant. The use of calcium dyes has been previously applied to the whole heart to measure calcium handling of specific regions of the heart [6-9]. The limitations of these experimental approaches are long incubation times *ex-vivo* to load the dye, uneven loading, and motion artefacts if single wavelength dyes are used.

Recently Dong et al., developed a ratiometric calcium reporter mouse called Salsa6f which expresses the genetic cytosolic calcium indicator gCaMP6F fused to tdTomato upon expression of cre [10]. In contrast to a dye-based approach, gCaMP and its derivatives use a M-13 calmodulin peptide to bind and buffer calcium. Genetic calcium indicators have been extensively used in *in-vitro* settings and in neuroscience applications [11, 12]. However, genetically designed calcium indicators have not been widely used for *in-vivo* cardiac research, with early reports of dilated cardiomyopathy with cardiac expressing GFP-mice [13]. The combination of genetically expressed calcium indicators with transgenic mice to examine genes implicated in cardiac disease processes is a powerful application if the limitations of GECIs in cardiomyocytes can be addressed. Indeed, it has been recently reported that gCaMP6F can be expressed in adult cardiomyocytes for *in-vivo* settings [14]. Yet, direct comparisons measuring gCaMP6F calcium transients in isolated cardiomyocytes to the existing Fluo4-AM literature have not been made.

Here, we tested the ability of the Salsa6f calcium reporter mouse to measure calcium transients in cardiomyocytes. We addressed the cardiotoxic effects of GFP and calcium buffering, by using an inducible αMHC-Mer-Cre-Mer mouse to enable temporally induced expression of Salsa6f in adult cardiomyocytes. We compared the cytosolic calcium transients of mice expressing Salsa6f to cardiomyocytes loaded with the calcium dye Fluo-4AM and recorded spontaneous calcium transients in the right atria as a proof-of-concept demonstration to measure atrial calcium handling. To assess feasibility of using the Salsa6f in transgenic mouse lines targeting genes that modify calcium handling, we crossed the Salsa6f mouse with a floxed *Pkd2* mouse. The gene *Pkd2* encodes the protein Polycystin-2, a non-selective cation TRP channel that localizes to the SR where it can mediate calcium transients via regulation of the ryanodine receptor and the L-type calcium channel [15]. Application of the Salsa6f model in a heterozygous and homozygous mouse model of cardiac PC2-KO demonstrates the feasibility of using the Salsa6f to measure cardiac calcium transients in transgenic models.

## 2. Materials and Methods

### Animal model

gCaMP6F-tdTomato expressing mice [10], inserted in the Rosa26 locus (Jackson Laboratories), were crossed with αMHC Mer-Cre-Mer (Jackson laboratories) to obtain gCaMP6F-tdTomato-cardiomyocyte specific mice under tamoxifen induction, hereafter called Salsa6f mouse. Some mice were further crossed with *Pkd2* floxed mice to generate gCaMP-tdTomato-Pkd2F and gCaMP-tdTomato-Pkd2FF mice. All studied mice had one copy of gCaMP6F-tdTomato. At 6 weeks of age, mice were given tamoxifen chow (tamoxifen citrate, 250mg, 10 days, 5 days on, 2 days off) as previously described ([16]), to induce Cre expression. Male and female mice, 13-20 weeks of age (1.5-3 months post tamoxifen induction) were used in the subsequent experiments. All animal studies performed were done under approved Institutional Animal Care and Use Committee (IACUC) protocols at Loyola University Chicago.

### Echocardiography

Mice (12-16 weeks of age; 1.5-2 months post tamoxifen) were anesthetized with isoflurane (2%) and kept at 36°C during the procedure. Hearts were imaged using the Vevo3100 (FUJIFILM Visual Sonics) using the M-mode (short axis). Left ventricular cardiac parameters were analyzed with VevoLab software from the short axis.

### Isolation of whole hearts and single cell cardiomyocyte

The excision of the whole heart for whole heart or single cell cardiomyocyte isolation was performed as previously described [16]. Mouse hearts were rapidly excised following anesthetization with isoflurane. The hearts were transferred to ice-cold perfusion buffer (in mM: 113 NaCl, 4.7 KCl; 0.6 KH_2_PO_4_; 0.6 Na_2_HPO_4_; 1.2 MgSO_4_; 12 NaHCO_3_; 10 KHCO_3_; 10 HEPES; 30 Taurine; pH 7.3), and canulated through the aorta. After cannulation, the hearts were perfused through the coronary arteries for 2-3 minutes with perfusion buffer. Whole hearts were then connected to a Variable Flow Peristaltic Pump (Thermo Fisher Scientific) in a custom-built platform for imaging (atrial imaging) or digested into individual cardiomyocytes with liberase as previously described [16].

### Whole heart imaging

Isolated whole hearts were reverse perfused through the aorta at a rate of 2 ml/min heated to 37°C with a final bath temperature of 34^0^C with a TC-324C Temperature Controller (Warner Instruments) and imaged through a 35 mm glass bottom plate (CellVis, California). Whole hearts were first perfused with perfusion buffer (in mM: 113 NaCl, 4.7 KCl, 0.6 KH_2_PO_4_, 0.6 Na_2_HPO_4_, 1.2 MgSO_4_, 12 NaHCO_3_, 10 KHCO_3_, 10 HEPES, 30 Taurine. pH 7.35) for 2 minutes and switched to Tyrode’s solution (in mM: 140 NaCl, 4 KCl, 2 MgCl2, 5 EGTA, 10 glucose, 10 HEPES, pH 7.4) with 25 μM of (S)-Blebbistatin (Cayman Chemicals) to reduce motion artefacts. After establishing a baseline of fluorescence for both gCaMP6F and tdTomato, with Tyrode’s solution, the beta-adrenergic receptor agonist isoproterenol (10 nM, Cayman Chemicals) was introduced to the perfusing solution for a total of 3 mins and the right atria was imaged. After stable chronotropic responses were achieved, the isoproterenol was washed out by returning the perfusate to Tyrode’s solution. gCaMP6F was excited with a 488nm LED while tdTomato was excited with 540 nM LED (Lumencor Spectra X lamp). Images of the right atria were acquired every 20 ms with a sCMOS camera (Orca Flash, Hamamatsu) with a 2.5x objective on the Zeiss Axio Observer 7 inverted fluorescent microscope using the Zen Blue software (Zeiss, Germany). The total time of cardiac imaging was kept to 1 hour post excision.

### Image analysis

After image acquisition, videos were analyzed through FIJI [17]. Videos were processed with image registration using the Rigid registration plugin. Five regions of interest were selected in the right atria and analyzed through the different conditions using the Time Series Analyzer V3 plugin.

### Measurement of [Ca^2+^] in cardiomyocytes with fluo4AM and comparison with Salsa6f

Cardiomyocytes from mice age-matched to the Salsa6f mice were isolated as described above and loaded with Fluo-4AM as previously described [16]. In brief, cells were allowed to settle on laminin coated coverslips and line scan confocal microscopy mode (Biorad) used to monitor calcium transients with a 488 nm excitation laser. The same experimental paradigms were conducted between Salsa6f and Fluo-4AM loaded cells, with cells subjected to the same pacing frequency, the same caffeine protocol and same isoproterenol conditions. To calculate the transient amplitude, the amplitudes of four voltage evoked responses were averaged. To calculate tau, a single exponential decay was fitted to the average of four voltage evoked responses. For measurements of tau with isoproterenol in paired cells, the calculation of tau was bounded by the maximum amplitude seen without isoproterenol. This was to ensure that the uptake measurement of SERCA with and without isoproterenol was calculated with the same calcium load.

### Statistical analysis

Data was plotted using GraphPad PRISM 9. For comparison of two independent groups statistical significance was analyzed through student’s t-test where Gaussian distribution was not assumed. Comparison of multiple groups were analyzed through One-Way ANOVA followed by Tukey’s multiple comparison test. Where appropriate, correlation tests were performed to compare different conditions within the same group. Conditions were considered statistically significant when p values were <0.05. Error bars indicate Standard Error of the Mean (SEM).

## 3. Results

### 3.1 Inducible expression of Salsa6f does not affect cardiac function

Long-term expression of fluorophores in cardiac applications have not been widely embraced due to early reports that the expression of green fluorescent proteins (GFPs) results in dilated cardiomyopathy [13]. Therefore, we first assessed that expression of the ratiometric GECI did not affect cardiac function through the echocardiography of Salsa6f mice in comparison to non-expressing Salsa6f mice (called control mice) 6-8 weeks after tamoxifen induction (Fig. 1). Analysis of the left ventricle mass/BW, ejection fraction, and fractional shortening remained unchanged between the control and the Salsa6f groups, and there were no differences between male or female mice (Fig. 1B-D). As dilated cardiomyopathy had been previously noted, we carefully measured the chamber size in the M-mode by examining the inner diameter under systole and diastole (Fig. 1E), the posterior wall thickness (Fig. 1F), and the anterior wall thickness (Fig. 1G). In all these measurements there was no difference, except for a reduction in diastolic diameter in the males, but no change under systole. The mice were subjected to similar levels of anesthesia during the echocardiographic analysis, as demonstrated by similar heart rates across all groups (Fig. 1H). This data demonstrates that the expression of cardiac specific Salsa6f largely does not affect systolic or diastolic cardiac function or LV structure/architecture.

**Figure 1.**
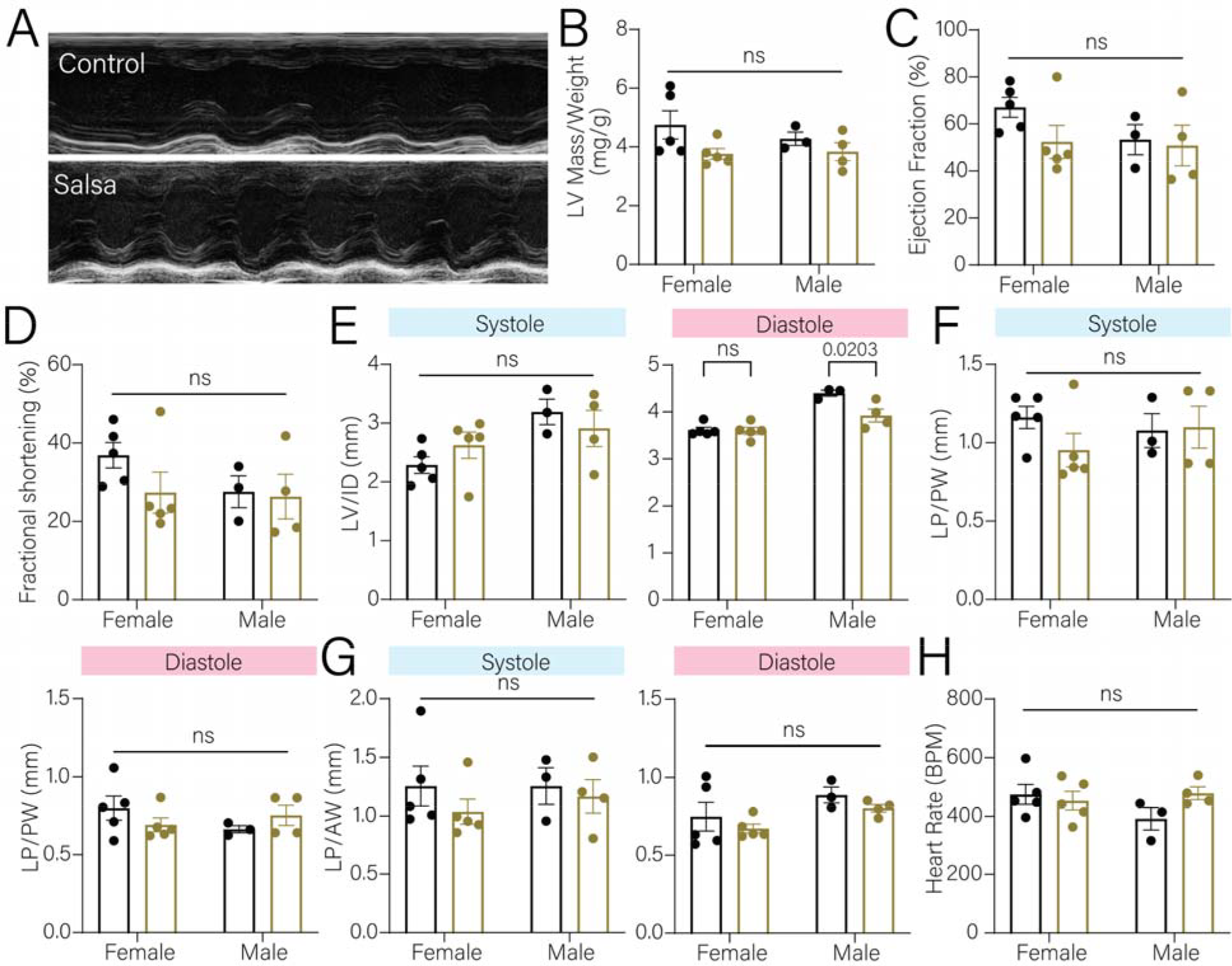
Cardiac function of Salsa6f expressing mice is not affected. **A**. Electrocardiogram of non-expressing Salsa6f (*Control*) mice (*top*) and Salsa6f expressing mice (*bottom*). **B**. No significant difference was observed of the left ventricle mass between the control (*black*) and the Salsa6f (*gold*) expressing mice. Dots represent individual mice. Bars represent mean±SEM. **C-D**. Quantification of the ejection fraction and fractional shortening showed no significant difference between control and Salsa expressing mice. Dots represent individual mice. Bars represent mean±SEM. **E-G**. Left ventricular diameter (LV/ID) (**E**), left ventricular posterior wall (LV/PW) (**F**), left ventricular anterior wall (LV/AW) (**G**) showed no difference between control and Salsa6f expressing mice. Dots represent individual mice. Bars represent mean±SEM. **H**. Heartbeat between the control and Salsa6f expressing mice are within normal range. Dots represent individual mice. Bars represent mean±SEM.

### 3.2 Imaging of single cell cardiomyocytes

We then examined the localization of the expression of both gCaMP6F and tdTomato in isolated cardiomyocytes. The expression of gCaMP6F and tdTomato was evenly distributed throughout the cytosol in the isolated single cardiomyocyte (Fig. 2A). Cardiomyocytes were electrically paced at 0.5 Hz under a wide-field fluorescence microscope and compared to the responses following the addition of isoproternol. Isoproterenol is a beta-adrenergic receptor agonist that has ionotropic effects due to the pleotropic effects of PKA phosphorylating the L-type calcium channel, the ryanodine receptor and lusitropic effects via phosphorylation of the SERCA regulator phospholamban [18]. As expected, isoproterenol amplified the calcium transients (Fig. 2B, *black vs blue*), with the systolic calcium amplitude significantly increased by 2-fold higher (Fig. 2C).

**Figure 2.**
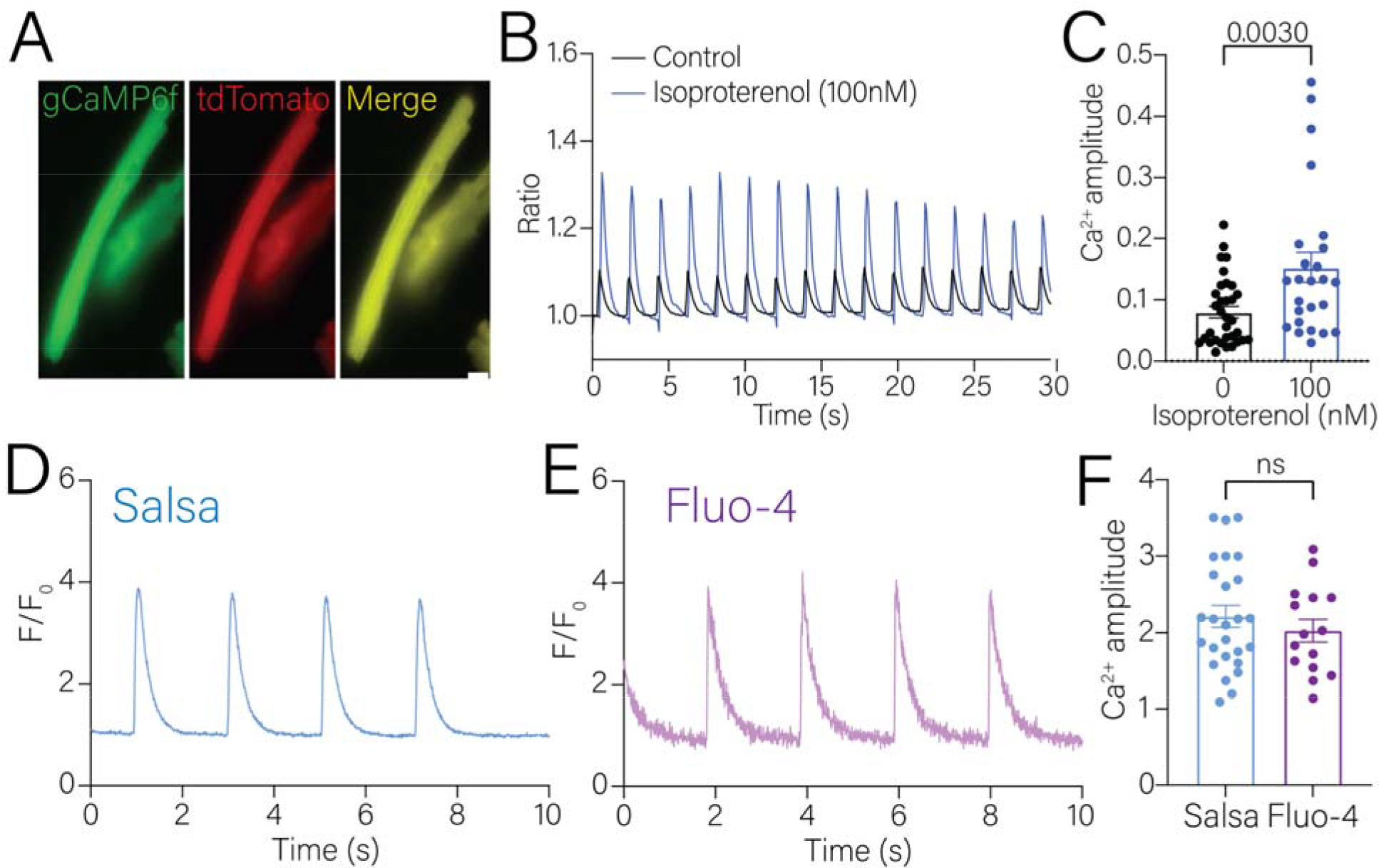
Measurement of calcium transients of isolated single cardiomyocytes. **A**. Representative images of isolated Salsa6f expressing cardiomyocytes. **B**. Representative calcium transients of electrically paced cardiomyocytes in the absence (*black line*) or presence of 10nM isoproterenol (*blue line*) acquired with wide-field microscopy. **C**. Cytosolic systolic calcium amplitude was significantly increased in isolated single cardiomyocytes in the presence of 10nM isoproterenol (*blue bar*). **D, E**. Representative calcium transients of Salsa expressing (**D**) or Fluo-4AM loaded (**E**) cardiomyocytes acquired with confocal line-scan microscopy. **F**. Calcium amplitude was similar in the electrically paced Salsa6f or Fluo-4AM loaded cardiomyocyte. Dots represent individual cardiomyocytes analyzed. Statistical analysis was performed using student’s t-test using non-parametric follow up analysis.

To compare the Salsa6f signal to calcium dyes, we shifted to using line-scan confocal microscopy (Fig. 2D-E). We selected Fluo-4AM to compare against Salsa6f due to the similarity in the kinetics of calcium detection (K_d_ ∼345nM vs 375nM) and same wavelength excitation. Therefore, the experimental parameters established between each condition could be compared. In this series of experiments, we did not measure the tdTomato signal. The average maximal change in amplitude of the calcium transients in the Salsa6f cardiomyocytes with 0.5 Hz pacing was around 2.2, with an upper maximum of ∼3.5 (Fig. 2F, *blue bar*). Cells loaded with Fluo-4AM had an average maximal amplitude of 2, with an upper maximum of 3 (Fig. 2F, *purple bar*). Thus, Salsa6f and Fluo-4AM had similarly sized evoked calcium transient signals (Fig. 2F).

We then used an experimental protocol where we could evaluate multiple components of the calcium transient cycle in paired cells before and after isoproterenol (Fig. 3A, F). In these paired measurements, application of isoproterenol resulted in doubling of the amplitude (taken as a measure of the L-type and RyR contribution) in the Salsa6f cells (Fig. 3B). In Fluo-4AM cells there was a significant increase in the calcium transient amplitude, but the fold change was less than that seen with Salsa6f (∼1.5-fold change, Fig. 3G). The tau of decay was calculated to provide an estimate of the function of the calcium uptake pump, SERCA. In Salsa6f cells, the tau of decay of the calcium transients was ∼270 ms and decreased to ∼245 ms (Fig. 3C). In the Fluo-4AM cells, the tau of decay was longer (∼290ms) and decreased to ∼120ms after isoproterenol (Fig. 3H).

**Figure 3.**
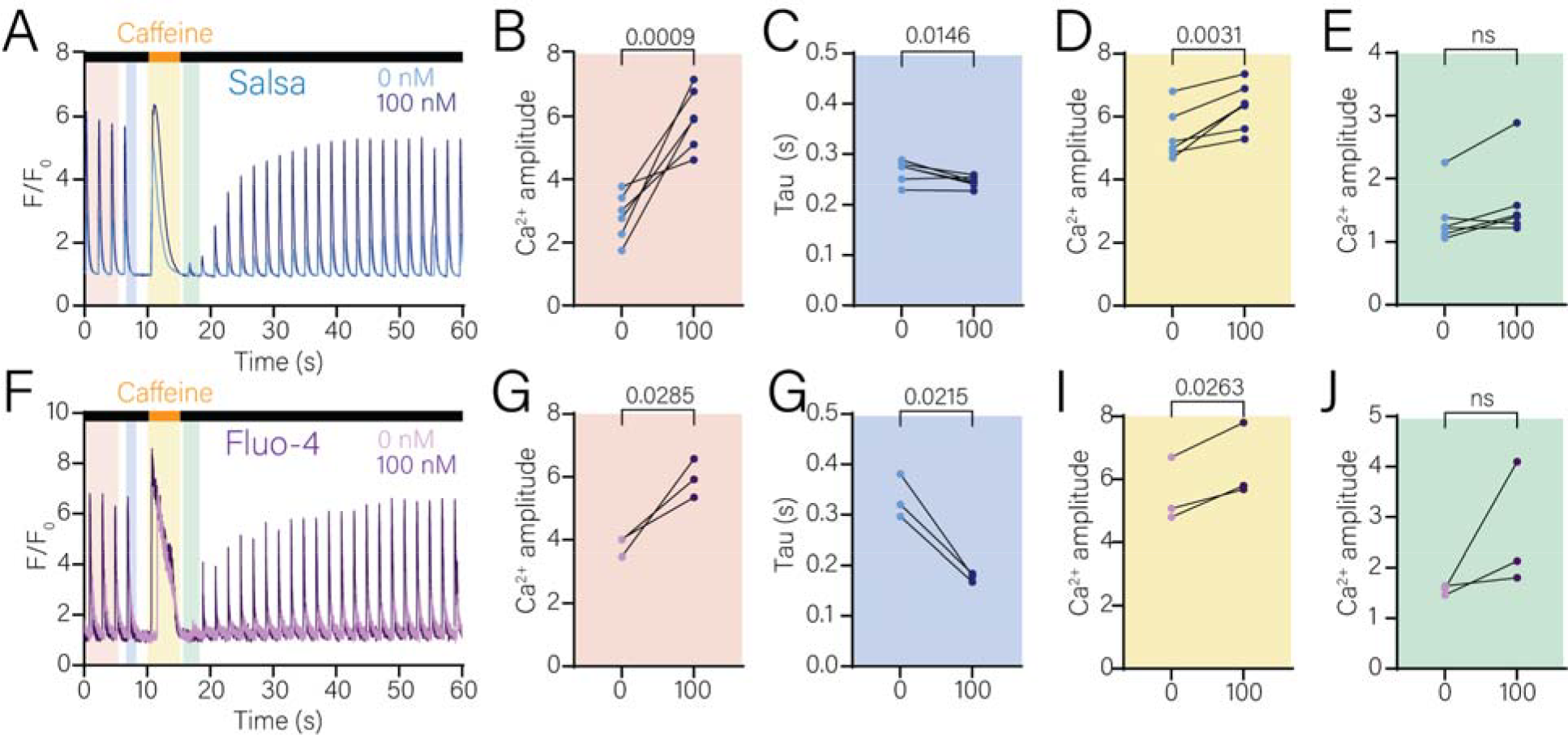
Paired measurements of calcium transients in cardiomyocytes. **A, F**. Representative calcium transients with a pacing-Caffeine(10mM)-pacing protocol in Salsa6f expressing (**A**) or Fluo-4AM loaded (**F**) cardiomyocytes before and after the addition of 100nM of isoproterenol. **B, G**. The voltage evoked calcium amplitude before and after presence of 100nM isoproterenol in Salsa6f expressing (**B**) or Fluo-4AM loaded (**G**) cardiomyocytes. **C, H**. The tau of decay of the evoked calcium amplitude before and after presence of 100nM isoproterenol in Salsa6f expressing (**C**) or Fluo-4AM loaded (**H**) cardiomyocytes. **D, I**. The caffeine evoked calcium amplitude before and after presence of 100nM isoproterenol in Salsa6f expressing (**D**) or Fluo-4AM loaded (**I**) cardiomyocytes. **E, J**. The calcium amplitude of the first evoked transient after store depletion, representing the L-type current, before and after presence of 100nM isoproterenol in Salsa6f expressing (**E**) or Fluo-4AM loaded (**J**) cardiomyocytes Dots represent individual cardiomyocytes analyzed. Statistical analysis was performed using paired analysis and student’s t-test. p-values are listed in the figure.

Pacing was ceased, and the SR load was depleted by the addition of caffeine (10mM) (Fig. 3A, F). In Salsa6f cells, the caffeine evoked amplitude was higher than the paced amplitude and was higher still with isoproterenol (Fig. 3D). In the Fluo-4AM cells, a similar response was observed (Fig. 3I). To examine the L-type calcium channel signal, the caffeine was removed, and the pacing protocol resumed. The first electrically induced calcium transient after store depletion was taken to be indicative of the L-type voltage-activated calcium channel. In Salsa6f cells, this amplitude was around 1.1-1.3 of the evoked amplitude, ∼10-25% of the original voltage evoked amplitude and did not change with isoproterenol application (Fig. 3E). In Fluo-4AM loaded cells, the L-type current amplitude was around 1.3-1.5 of the evoked amplitude, ∼25%-30% of the original voltage evoked amplitude (Fig. 3J). There was a trend upwards in the amplitude of the signal after isoproterenol application, but it did not reach significance.

Overall, there was a similar dynamic range in the calcium signals evoked by voltage between Salsa6f and Fluo-4AM. However, there was a greater fold change in the amplitude of the signal following addition of isoproterenol in the Salsa6f cells compared with Fluo-4AM cells. The tau of decay was faster in Salsa6f cells compared to the Fluo-4AM cells. This result may likely reflect the faster off kinetics of calcium unbinding from Salsa6f than the unbinding of PLB to SERCA. It should also be noted that from the Salsa6f cells analyzed, the L-type transient was barely detectable in ∼10% of all cells, suggesting that there could be buffering of the L-type signal due to the calmodulin binding domain of the gCaMP6F signal interfering with calcium handling molecules such as L-type voltage gated calcium channels and calcium-binding proteins [19].

### 3.3 Isolation and imaging of Salsa6f in *ex-vivo* hearts

One potential advantage of a genetically encoded indicator is the ability to measure calcium transients in the whole heart without lengthy dye incubation times and uneven loading. Additionally, our isolated cardiomyocyte experiments indicated that the signal-to-noise ratio would be favorable. To test if the Salsa6f mouse could be used for this purpose, we excised and cannulated the heart via the aorta in a Langendorff preparation. Once cannulated, the heart was put in a glass bottom dish for imaging and connected to a custom-made platform where the heart was attached to a peristaltic pump (Fig. 4A). We imaged different areas of the heart (atria and ventricles) where we observed evenly distributed cytosolic expression of both gCaMP6F and tdTomato (Fig. 4B). The expression of Salsa6f was not observed in blood vessels or other non-cardiac tissue. To minimize artifacts caused by motion of the spontaneous beating heart, the hearts were perfused with Tyrode’s solution that contained 25 μM of (S)-Blebbistatin, which inhibits myosin. Once stability was achieved, we recorded calcium transients and measured an average of 102 beats per minute in the atria, and no evidence of arrhythmogenic activity (Fig. 4C, *baseline*). To demonstrate that the excised heart can be pharmacologically stimulated we perfused the hearts with isoproterenol. Following the addition of isoproterenol, we observed an increase to an average of 177 beats per minute in comparison to the baseline (Fig. 4C, *isoproterenol*). Quantification of the calcium amplitude before and after the addition of isoproterenol showed a significant difference in the calcium amplitude (Fig. 4D). Taken together, we demonstrate that we can detect atrial calcium transients through the expression of Salsa6f in an intrinsic beating heart and stimulate the heart through pharmacological approaches.

**Figure 4.**
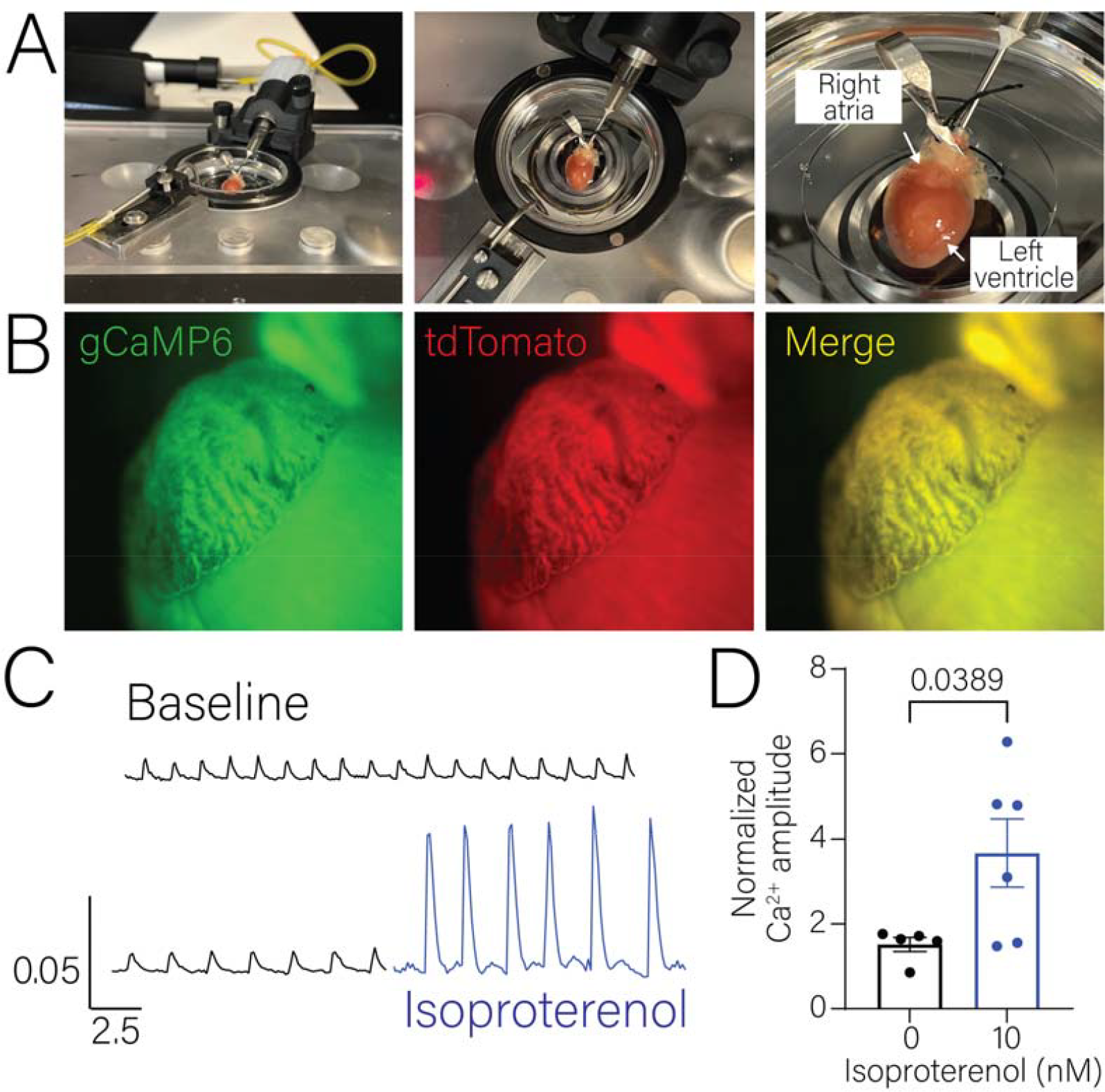
Imaging of excised whole hearts expressing Salsa6f. **A**. Images of the experimental and imaging setup of the excised hearts expressing Salsa6f. **B**. Representative images of Salsa6f expressing right atria. **C**. Representative tracing of baseline calcium transients (*top*) and after the addition of 10nM of isoproterenol (*bottom*) of Salsa6f transient hearts. **D**. Systolic calcium amplitude was significantly increase after the addition of 10nM of isoproterenol. Dots represent individual mice. n=5-6 mice. Bar graphs represent mean±SEM. Statistical analysis was performed using student’s t-test. p-values are listed in the figure.

### 3.4 Deletion of cardiomyocyte localized Polycystin-2 increases calcium release

A practical application of the Salsa6f mouse is in combination with floxed mice targeting genes that regulate cardiac calcium signaling. As proof-of-principle, we crossed the Salsa6f mouse with Polycystin-2 (*Pkd2*) floxed mice to create inducible heterozygous or homozygous deletion of cardiac Polycystin-2. Polycystin-2 (PC2) is a non-selective cation channel from the Transient Receptor Potential (TRP) Family, in which loss-of-function mutations can lead to the development of ADPKD where the main cause of mortality is cardiac failure [20]. Quantification of the ex-vivo hearts averaged 141 beats per minute in the heterozygous mice versus an average of 112 in the homozygous PC2 knock-out mice (Fig. 5A-B). The addition of isoproterenol in the PC2 cardiac knockout significantly increased the heartbeat, which was not observed in the heterozygous knock-out mice (Fig. 5B).

**Figure 5.**
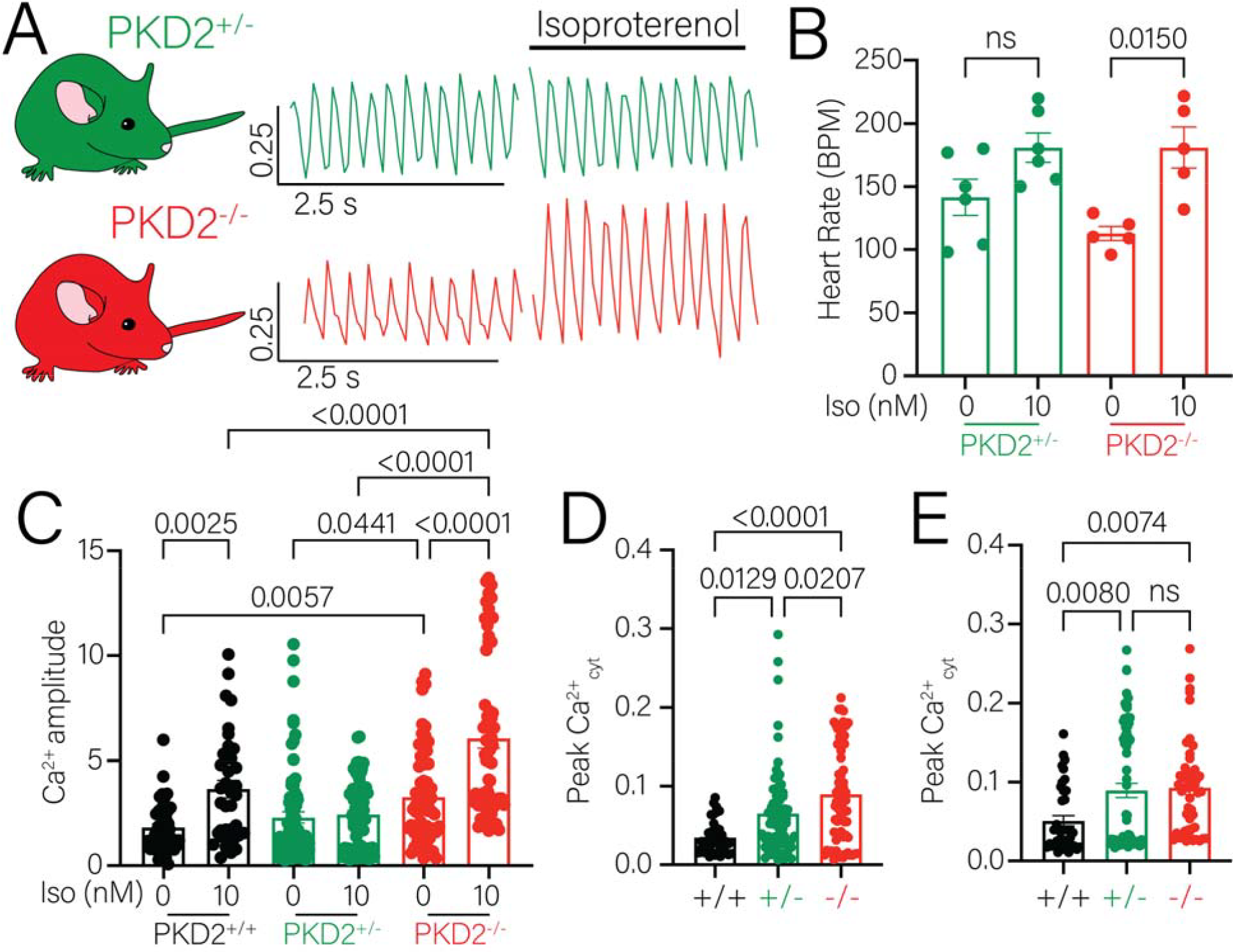
Analysis of calcium transient from whole hearts in heterozygous and homozygous cardiac deletion of PC2 mice. **A**. Representative calcium transient before and after addition of isoproterenol (10nM) in PC2 heterozygous (*top*) or knock out (*bottom*) mice. **B**. Heart rate of heterozygous mice was not increased after the addition of isoproterenol while it significantly increased in the knockout mice. Bar graphs are mean±SEM. Dots represent individual mice (n=5-6 mice). Statistical analysis was performed using 2-way ANOVA. p-values listed in figure. **C**. Calcium amplitude comparison between control (*black bars*), PC2 heterozygous (*green bars*) and PC2 knock-out (*red*) in the absence or presence of isoproterenol. Bar graphs are mean±SEM. Dots represent quantification of 5 sections in the right atria from 5-6 mice. Statistical analysis was performed using 2-way ANOVA. p-values listed in figure. **D**. Peak cytosolic systolic calcium was significantly increased in PC2 heterozygous (*green bars*) and PC2 knock out (*red bars*). Statistical analysis was performed using One-way ANOVA. p-values listed in figure. **E**. Peak cytosolic systolic calcium was significantly increased in the presence of isoproterenol in both PC2 heterozygous (*green bars*) and PC2 knock out (*red bars*). Bar graphs are mean±SEM. Dots represent quantification of 5 sections in the right atria from 5-6 mice. Statistical analysis was performed using One-way ANOVA. p-values listed in figure.

As shown in control mice (Fig. 4), addition of isoproterenol significantly increased cytosolic calcium (Fig. 5C, *black bars*). Prior to isoproterenol addition, no difference was observed of systolic calcium levels in heterozygous knock out mice in comparison to the control mice but significantly increased in homozygous knock out mice (Fig. 5C, *compare 0 bar graphs*). Previous work demonstrated that cardiac PC2 increases SR calcium release by modulating the RyR activity [16], which is congruent with increased systolic calcium levels. Addition of isoproterenol in both control and homozygous knock out PC2 mice significantly increased cytosolic calcium. However, cytosolic calcium did not increase in the heterozygous PC2 mice. This is consistent with a lack of increase in the heartbeat after the addition of isoproterenol. Quantification of peak cytosolic calcium was significantly increased in hetero- and homozygous PC2 knock out mice during systolic levels (Fig. 5D) and after the addition of Iso (Fig. 5E). Taken together this data demonstrates the feasibility of Salsa6f to study calcium transients in a cardiac specific application. Additionally, we show that deletion of cardiac PC2 leads to an increase in cytosolic calcium which is not observed in a heterozygous model.

## 4. Discussion

Cardiac contractility requires the increase of cytosolic calcium due to both calcium influx across the plasma membrane and calcium release from the SR, and the relaxation phase requiring calcium uptake by SR and plasma membrane pumps and exchangers. Cardiac calcium dysregulation is the basis of several diseases that lead to arrythmias, cardiac dysfunction and failure [21, 22]. Thus, the ability to measure calcium signals in isolated cardiomyocytes and whole heart preparations reliably without perturbation of cardiac function is essential. In this study we present the adaptation of the ratiometric genetically encoded calcium indicator Salsa6f in an inducible mice model to study cytosolic calcium transients in the heart. We demonstrate that the inducible expression of the Salsa6f does not affect the gross anatomical or function of the heart as assessed by echocardiography. We showed the ability to measure calcium signals in isolated single cardiomyocyte and spontaneous calcium transients in an intact atrial preparation. Lastly, we demonstrate the feasibility of this system when combined with heterozygous and homozygous models of cardiac specific PC2 KO.

In recent years, the application of genetically encoded calcium indicators (GECIs) has broadened our understanding of calcium handling in a variety of different systems. However, the application of these GECIs in cardiac settings is limited. This has in part been due to early studies demonstrating that the expression of GFP in cardiomyocytes from birth caused dilated cardiomyopathy, most likely due to the interaction of GFP with the myofilament [13]. To overcome the cardiotoxicity of GFP, we therefore used an inducible cardiac model to temporally restrict expression. In contrast to the previously reported dilated cardiomyopathy, we did not note any major structural remodeling with echocardiography up to 3 months after Salsa6f expression. It has also been noted that tamoxifen can similarly induce cardiac remodeling [23]. We therefore used a dietary formulation of tamoxifen to minimize any cardiac effects [16].

Most measurements of calcium in cardiomyocytes use dye-based approaches. There are differences in the dynamic range and calcium sensing capabilities of GECIs in comparison to calcium indicators like Fluo-4AM. The calcium K_d_ between gCaMP6F and Fluo-4AM are relatively similar (∼345nM vs 375nM) [3, 4]. However, on-GECIs calcium binds to the calmodulin binding region within the protein causing a conformational change which leads to an increase of fluorescence [3]. In contrast, Fluo-4AM is a chemical dye based on calcium chelation which causes a spectral shift [24]. Despite these different binding properties, we have found that the long-term expression of Salsa6f in cardiomyocytes can faithfully report calcium transients with similar values to Fluo-4AM. Importantly, cardiomyocytes expressing Salsa6f have a similar amplitude range to cardiomyocytes loaded with Fluo-4AM and have a robust positive response to ionotropic stimuli (isoproterenol).

We note two areas of caution with cardiac applications of Salsa6f. Calmodulin is known to bind and/or regulate calcium channels like the L-voltage gated calcium channels, RyR and SERCA [25-27]. In >80% of cells expressing Salsa6f, we found that the L-type calcium signal could be measured and was of a similar amplitude to that measured with Fluo-4AM. However, in ∼10% of cells, there was a reduction in the ability of the Salsa6f expressing cardiomyocytes to detect the voltage activated calcium entry after depletion of the SR, which is likely due to the binding of the calmodulin region of the gCaMP6F to these proteins, as has been observed in neuronal applications [19]. The possibility of gCaMP6F binding to different proteins within the cell and the buffering capacity should be taken into consideration when interpreting the data. We also noted that the tau of decay for the evoked calcium transients was faster in Salsa6f cells compared with Fluo-4AM or with values calculated with Fura-2AM [16]. This is likely due to the fast off-kinetics for gCaMP6F compared with Fluo-4AM and Fura-2AM [28]. Although the tau was significantly faster in both Salsa6f and Fluo-4AM cells following isoproterenol addition, the change was not as large in the Salsa6f mice, suggesting there may be subtle changes in calcium handling occurring with long-term expression of Salsa6f including chronic phosphorylation of PLB, or downregulation of PLB. However, when these data are taken together with the overall findings from our analysis of the cardiomyocyte calcium transient suggests that Salsa6f does not interfere with the calcium handling machinery.

## 5. Conclusion

We have demonstrated that Salsa6f transgenic mouse model can be used to study calcium transients from the single cardiomyocyte cell to the whole heart to uncover and understand how individual cellular processes result in whole organ physiological processes. The use of the Salsa6f mouse in cardiac applications has some limitations, but importantly, there is little cardiotoxicity with this animal model up to 3 months after induction. Calcium transients from cardiomyocytes isolated from these mice show similar properties to cells loaded with the dye Fluo-4AM. Moreover, these mice display appropriate positive responses to ionotropic agonist stimulation. We have also shown that the Salsa6f mouse can be used to interrogate atrial calcium handling, and specifically applied it in the context of studying the Polycystin-2 protein in the heart. We therefore contend that the Salsa6f mouse, when used carefully, is an aid to cardiac researchers to investigate calcium handling from the single cell to the whole heart and in conditions that lead to cardiac failure.

## Acknowledgements

We thank the Kuo lab members and Dr. Seth Robia (Loyola University Chicago) for helpful discussions. Grants and funding: R00DK101585 (IYK); R01HL151990 (AZ). The Loyola University Chicago CVRI acknowledges NIH grant 1S10OD028449 to support purchase of the Vevo3100 ultrasound Research reported in this publication was supported by the National Institute of Diabetes and Digestive and Kidney Diseases of the National Institutes of Health under Award Numbers U2CDK129917 and TL1DK132769 (KMMN).

## Contributions

Conception: KMMN and IYK. Experiments: KMMN, EB, QC and IYK. Analysis: KMMN, and IYK. Discussions: KMMN, EB, AZ and IYK. Draft: KMM and IYK. All authors approved the final manuscript.

